# Angiotensin II Infusion Promotes Activation and Selective Cytokine Responses in Activated CD4 and CD8 T cells

**DOI:** 10.1101/2025.10.01.679792

**Authors:** Devon A. Dattmore, Jeffrey R. Leipprandt, Saamera Awali, McKenzie Mahlmeister, D. Adam Lauver, Cheryl E. Rockwell

## Abstract

Hypertension affects roughly half of adults in the U.S. and is caused by several factors, including elevated angiotensin II (Ang II). T cells have been implicated in mechanisms of this pathology, however, data regarding T cell activation, including cytokine secretion and induction of activation markers are limited. This study investigated the hypothesis that Ang II increases T cell activation, as indicated by altered cytokine secretion and expression of surface markers associated with activation. To test this, 11-week-old C57Bl/6J male mice received saline vehicle or Ang II (490 ng/kg/min) via osmotic pump for 14 days (n=10/group) followed by immune cell isolation and ex vivo activation. Splenic T cells from Ang II-infused mice had modestly increased expression of CD25 (CD4: p=0.024, CD8: p=0.007), CD69 (CD4: p=0.017, CD8: p=0.032), and CD137 (CD8: p=0.022) 24 hours post-activation, as measured by median fluorescent intensity. The increased expression of activation markers correlated with an increase in select cytokines, including interleukin (IL)-28B (IFNλ3) and interferon (IFN)-γ-induced protein 10 (IP-10) at 24 hours and IFNγ and IP-10 at 120 hours, while decreasing IL-23 at 120 hours. The increased expression of CD25 and CD69 suggests Ang II may increase the magnitude of T cell activation. This is further supported by the elevated induction of select cytokines all of which are associated with an antiviral response. Taken together, the data suggest Ang II modestly promotes T cell activation resulting in selective induction of cytokines associated with antiviral immunity.

## Introduction

In the United States, roughly half adults have hypertension (blood pressure (BP) consistently higher than >130/80 mmHg). This is of concern, as recent reports indicate(1)(1)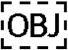. Uncontrolled hypertension results in progression of the condition, resulting in increased disease severity and complications, which portend increased mortality and health care costs. Hypertension is a complex disease, with several different etiologies. Disease progression, especially in the case of uncontrolled hypertension, however, is generally associated with chronic, low-grade inflammation. The mechanism(s) by which this inflammation occurs is not fully understood. Expanding our understanding of immune system dynamics in hypertension and its associated diseases, like metabolic syndrome, could lead to novel therapeutic approaches that improve patient outcomes.

In the past couple decades, there has been growing attention to the role that the immune system plays in hypertension, and metabolic syndrome, broadly. Hypertension and metabolic syndrome are frequently associated with an inflammatory immune profile that is characterized by in increases in pro-inflammatory M1 macrophages, double negative (IgM^-^ CD27^-^) B cells, and Th1 and Th17 T cells (2). The reason(s) for this phenotypic skewing is not fully understood, and there are multiple hypotheses for why this occurs. It is likely that this phenomenon is multi-factorial, with contributions from microbiotic changes in the gut, alterations in fatty acid profiles, vascular stress, neuronal inputs, alterations in immunomodulatory mediators, and more. Among the immunomodulatory mediators, Ang II is well established as a blood-pressure modulator and has also been shown to exhibit pro-inflammatory effects.

A meta-analysis of randomized controlled trials demonstrated that blocking the actions of Ang II, either through angiotensin-converting enzyme inhibitors (ACEi’s), or Ang II receptor blockers (ARB’s), has anti-inflammatory effects (decreased c-reactive protein, interleukin (IL)-6, and TNFα) (3). This is noteworthy because there is evidence that circulating IL-6 data may be predictive of hypertension onset (4). Additionally, studies in animals have shown anti-inflammatory effects, both *in vivo* and *in vitro*. Numerous studies indicate that Ang II promotes M1 skewing in monocytes/macrophages (5-7). There are only a handful of studies investigating Ang II effects on T cells, though the limited data suggest that Ang II exerts pro-inflammatory effects in T cells (8-12).

There are multiple lines of evidence that T cells play an essential role in hypertension. Mycophenolate mofetil (MMF), which diminishes lymphocyte proliferation, has been shown to lower blood pressure in hypertensive patients with psoriasis and rheumatoid arthritis (13). In the Sprague Dawley rat, MMF treatment prevented salt-sensitive hypertension following three weeks of Ang II infusion. The MMF treatment did not affect the hypertension in the first three weeks of the Ang II infusion (14). In 1976, Svendsen published their findings that mice with genetic aplasia of the thymus, deficient in mature T cells, were protected against DOCA-induced hypertension. Furthermore, thymus grafting into these mice removed the protective phenotype (15). Subsequent studies by Khraibi et al., revealed that T cells from hypertensive animals can elicit hypertensive phenotypes in non-hypertensive animals, demonstrating a phenotypic difference in T cells, which is capable of modulating blood pressure (16). Additionally, Dahl salt-sensitive (SS) Rag1^-/-^ rats, which have diminished lymphocytes, have reduced blood pressure and albuminuria on a high salt (4% NaCl) and high protein (30%) compared to SS counterparts (17). Rag1^-/-^ mice (C57BL/6 background) did not develop hypertension in response to a two-week Ang II infusion via osmotic pump (490ng/kg/min). What’s more, transplantation of T cells (but not B cells), was able to restore the hypertensive phenotype in these mice. Beyond this, transplantation of T cells from AT_1_R^-/-^ mice did not restore the hypertensive phenotype, suggesting that Ang II directly affects T cells through AT_1_R, leading to the development of hypertension. *In vitro* activation (anti-CD3) of peripheral blood T cells from Ang II infused mice resulted in higher IFNγ and TNFα production. *In vitro* exposure of splenocytes to Ang II (100nM), followed by activation (anti-CD3), resulted in higher supernatant levels of TNFα. Treatment with etanercept (8 mg/kg), an anti-TNFα biologic, decreased blood pressure in Ang II infused mice. Collectively, these data suggest that Ang II contributes to increased T cell activation and the resulting TNFα production contributes to development of hypertension (8).

Another study found that antigen-specific splenic CD8 T cells from spleens of AT_1_R^-/-^ mice have decreased CD25 MFI and are less likely to be positive for IL-2, CD25 (IL-2RA), and CD122 (IL-2B), 1 to 2 weeks post-immunization. Normally, following immunization, the percentages of CD69, CD44, PD1, LAG3, and CTLA4 positive antigen-specific splenic CD8 T cells are initially elevated, which then slowly decrease over time. However, in AT_1_R^-/-^ mice, these populations continue to increase. These data suggest that angiotensin II plays an important role in CD8 T cell activation and exhaustion (10). What’s more, T cells produce their own endogenous Ang II, which is capable of contributing to self-activation. Silva-Filho et al. (2011), demonstrated that endogenous Ang II contributes to splenic T cell activation in response to anti-CD3 *in vitro*, while exogenous supplementation of Ang II (100 mM) failed to elicit an increased response (as determined via percentages of CD25^+^ and CD69^+^ T cells) (PMID: 21641648). This lack of activation response to exogenous Ang II is in contrast to the Guzik (2007) study, which observed increased IFNγ and TNFα in splenic T cells treated with anti-CD3. Aside from the differences in the parameters that were assessed (cytokines verses surface activation markers), the apparent difference in T cell activation response to exogenous Ang II may be explained by mouse strain differences. The Guzik study used splenocytes from C57BL6/J mice, whereas the Silva-Filho study used splenocytes from BALB/c mice. These strains have different immunophenotypes; Th1 skewed verses Th2 skewed, respectively (18). Beyond these studies, the role that T cells play in hypertension is reviewed further by others (19).

Collectively, these studies suggest that T cells contribute to Ang II-induced increases in blood pressure. What is more, these studies also contribute evidence that Ang II promotes T cell activation, under various conditions. These data are limited, however, and additional studies would provide additional insight into the scope and mechanism. In this study, the effects of Ang II on T cell activation were further investigated, using an *in vivo* administration of Ang II via osmotic pump, followed by polyclonal T cell stimulation of full splenocyte fractions. This study was designed to determine the impact of 2-week Ang II infusion on subsequent responsiveness of isolated immune cells to polyclonal T cell activation as determined by induction of activation markers and cytokines secretion.

## Methods

### Animals

All animal protocols and procedures were approved by the Institutional Animal Care and Use Committee at Michigan State University and are in concurrence with the Guide for the Care and Use of Animals. 10-week-old male C57BL6/J mice were purchased from Jackson Laboratories (Bar Harbor, Maine). Animals were maintained on a 12-hour light-dark schedule and were provided with food and water *ad libitum*. At 12-to 13-weeks of age, these mice were surgically implanted with osmotic pumps containing either saline or Ang II (490 ng/mL/min). Immediately thereafter, carprofen (5 mg/kg) and enrofloxacin (5 mg/kg) were administered via subcutaneous injection. Mice were euthanized via cervical dislocation following 14 days of saline/Ang II infusion.

### Splenocyte isolation and T cell activation

Using sterile technique, spleens were excised, mashed with a syringe plunger, and the resultant single cell suspension was filtered through a 70 μm cell strainer. The filtrate was then centrifuged at 300 xg (4°C), the cell pellet was resuspended in 10 mL sterile RPMI. Following a second wash with RPMI, cells were cultured in 200 μL complete RPMI (RPMI 1640, 10% fetal bovine serum, 100 U/mL penicillin, 100 U/mL streptomycin, 25 mM HEPES, 1 mM sodium pyruvate, and 10 mM non-essential amino acids) with or without a polyclonal activation cocktail (1.5 μg/mL anti-CD3, 1.5 μg/mL anti-CD28, 1.5 μg/mL crosslinker) in 96-well plates at a cell density of 2 x 10^6^ cells (for 24-hour timepoint) or 0.5 x 10^6^ (for 120-hour timepoint). Cells were collected at 24-hours and immediately processed for flow cytometry. Supernatants were collected at 24- and 120-hours and immediately stored at -80°C for future cytokine/chemokine analysis.

### Flow cytometry

Cells were FACS-labeled with a T cell activation antibody panel (Table 1). Fluorescence was detected and analyzed using an Attune NxT flow cytometer (Thermo Fisher Science Inc., Waltham, MA), with the following settings; flow rate: 100 mL/minute, total volume: 180 μL, draw volume: 150 μL, rinse Cycles: 2, mix Cycles: 1.

**Table 1.**

| Channel | Target | Fluorochrome | Clone | Company | Catalog # | Dilution |
| --- | --- | --- | --- | --- | --- | --- |
| BL1 | CD3 | Alexa Fluor<sup>®</sup> 488 | 17A2 | Biolegend | 100210 | 1:100 |
| RL1 | FoxP3 | Alexa Fluor<sup>®</sup> 647 | MF-14 | Biolegend | 126408 | 1:100 |
| RL2 | CD69 | Alexa Fluor<sup>®</sup> 700 | H1.2F3 | Biolegend | 104539 | 1:100 |
| RL3 | CD4 | APC/Cyanine7 | GK1.5 | Biolegend | 100414 | 1:100 |
| VL1 | CD62L | Pacific Blue<sup>™</sup> | MEL-14 | Biolegend | 104424 | 1:100 |
| VL2 | Zombie | Fixable Aqua<sup>™</sup> Dye | N/A | Biolegend | 77143 | 1:100 |
| VL3 | CD134 (OX-40) | Brilliant Violet 605<sup>™</sup> | OX-40 | Biolegend | 119419 | 1:100 |
| VL4 | CD25 | Brilliant Violet 711<sup>™</sup> | PC61 | Biolegend | 102049 | 1:100 |
| YL2 | CD44 | PE/Dazzle<sup>™</sup> 594 | IM7 | Biolegend | 103056 | 1:100 |
| YL3 | CD8a | PE/Cy5.5 | 53-6.7 | Invitrogen | 35-0081-82 | 1:100 |
| YL4 | CD137 (4-1BB) | PE/Cy7 | 17B5 | Invitrogen | 25-1371-82 | 1:100 |

### Cytokine/Chemokine Analysis

Splenocyte supernatants were analyzed by Eve Technologies Inc. (Alberta, Canada) using the Mouse Discovery 32-plex (MD32) and Mouse Discover 12-plex Th17 (MD12-TH17) array panels.

### Statistical Analysis

All statistical analysis and plotting of graphs were performed using Prism 10 GraphPad software. Mann-Whitney tests were used when analyzing data with non-Gaussian distributions. Otherwise, data were analyzed via unpaired two-tailed t tests. Welch’s corrections were implemented for comparisons of unequal variances. Outliers were identified and removed using Grubbs’ outlier test.

## Results

### CD8 T cells are more sensitive to the stimulatory effects of Ang II on induction of cell surface activation markers

Previous studies have shown that angiotensin II can promote expression of cell surface receptors and cytokines associated with T cell activation and effector function in various in vitro and in vivo models. These pioneering studies are intriguing but lead to more questions about the scope of the effects of angiotensin II on T cell activation and effector function. In this study, we infused male C57Bl/6 with Ang II for 2 weeks prior to immune cell isolation from spleen. We then activated the cells with the T cell activator, anti-CD3/anti-CD28, for 24 – 120 h. As expected, anti-CD3/anti-CD28 stimulation resulted in a rapid increase in CD25, CD44, CD69, CD134 and CD137 and a concurrent decrease in CD62L (Fig. 1).

**Figure 1.**
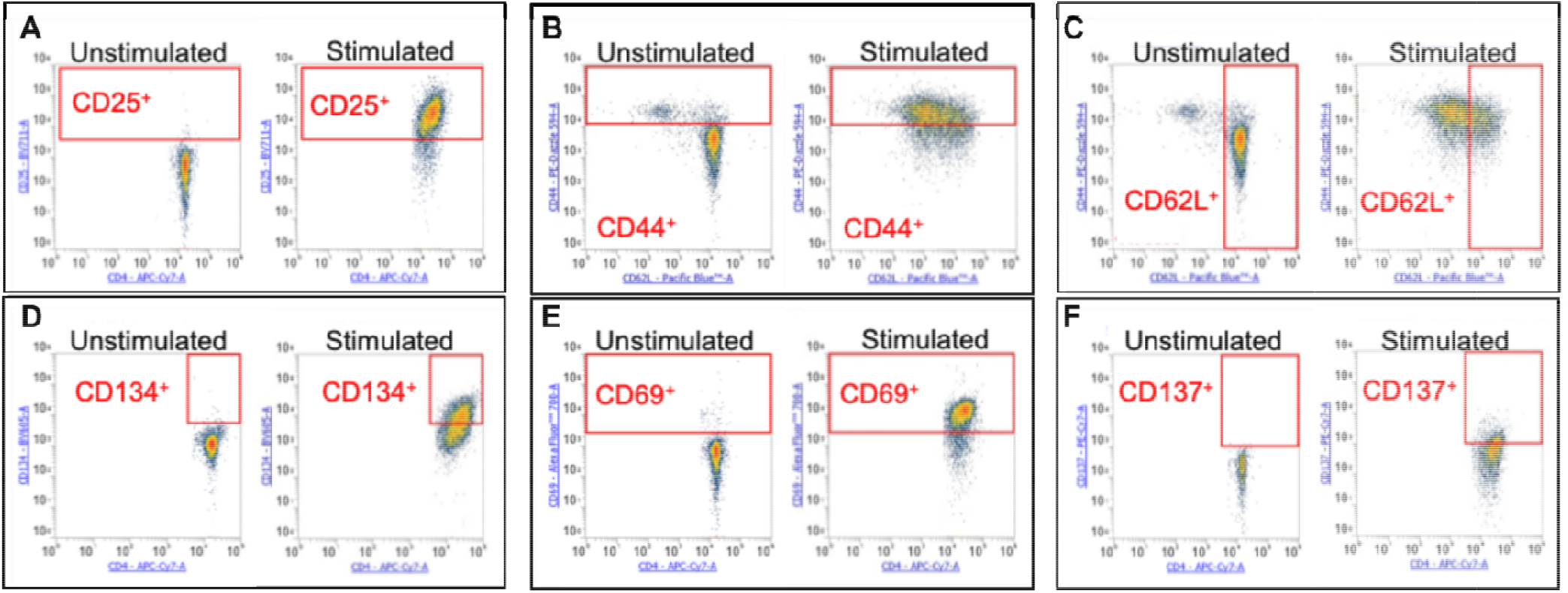
Effects of anti-CD3/anti-CD28 stimulation on T cell activation markers. Examples of T cell activation markers in CD4 ^+^ T cells 24 h post-stimulation (unstimulated vs stimulated). Live cells were gated using Zombie Aqua Dye from singlet populations (not shown). From the resulant live cells, CD4 and CD8 T cell populations were gated. Red boxes indicate gating strategy for each surface marker (all examples are CD4 ^+^ T cells; CD8 ^+^ T cell gating of surface activation markers was similar to CD4^+^ T cells and is not shown). A) CD25^+^ CD4 T cells, B) CD44 ^+^ CD4 T cells. C) CD62L^+^ CD4 T cells, D) CD69^+^ CD4 T cells, E) CD134^+^ CD4 T cells, F) D137^+^ CD4 T cells. Median fluorescent intensity (MFI) calculations were determined using the entire CD4^+^ or CD8^+^ populations.

Cellular markers of activation were increased in both activated CD4^+^ and CD8^+^ T cells from Ang II-infused mice, compared to saline-infused mice. In CD4^+^ T cells, CD25 and CD69 were increased, as determined by median fluorescent intensity (MFI) (Figure 2A). There was also a non-statistically significant trend toward an increase in CD134 by Ang II, which although similar in magnitude to the increases in CD69 and CD25 showed greater variability (Figure 2A). Similar, but more pronounced, findings were observed in CD8^+^ T cells, where Ang II increased expression of CD25, CD69, and CD137 as determined by MFI (Figure 2B). Interestingly, in contrast to CD69, CD25 and CD137, the activation markers CD44 and CD62L were unchanged in both CD4^+^ and CD8^+^ T cells (Figures 3A and 3B). Importantly, the percentage of T regulatory cells (Tregs) were also unchanged 24 hours post-stimulation and thus, the Ang II-mediated increases in activation markers cannot be attributed to a decline in the Treg population (Figure 3C). There were no differences between unstimulated T cells from saline and Ang II treated mice (data not shown). Collectively, these data indicate Ang II infusion promotes expression of the early activation markers CD25 and CD69 in both CD4 and CD8 T cells and this effect cannot be attributed to a decrease in the number of Treg’s.

**Figure 2.**
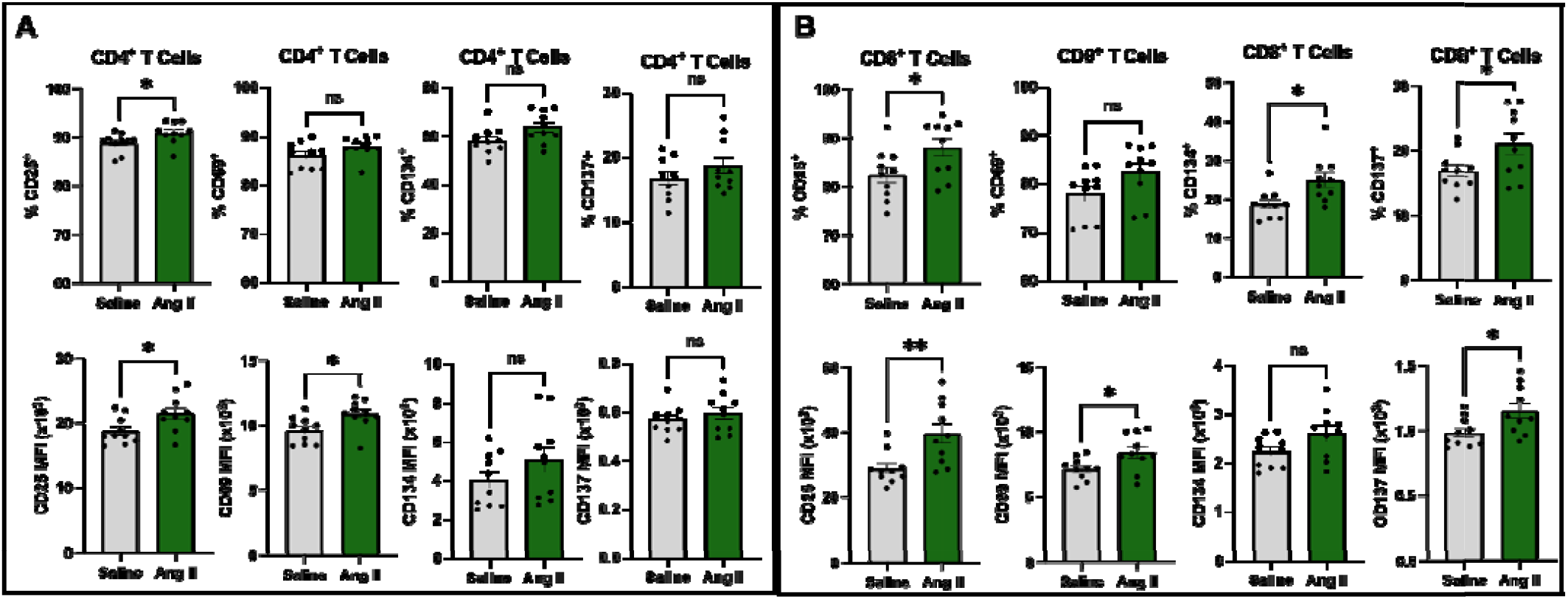
Effects of Ang II on T cell activation markers. T cell activation markers In A) CD4^+^ and B) CDS^+^ T cells) 24 h post-stimulation (n=9-10 mice/group). Results are depleted as mean ± SEM. Statistical analysis was conducted using unpaired two-tailed t-tests (Welch’s correction used for comparing data of unequal variances; *=p<0.05, **=p<0.01).

**Figure 3.**
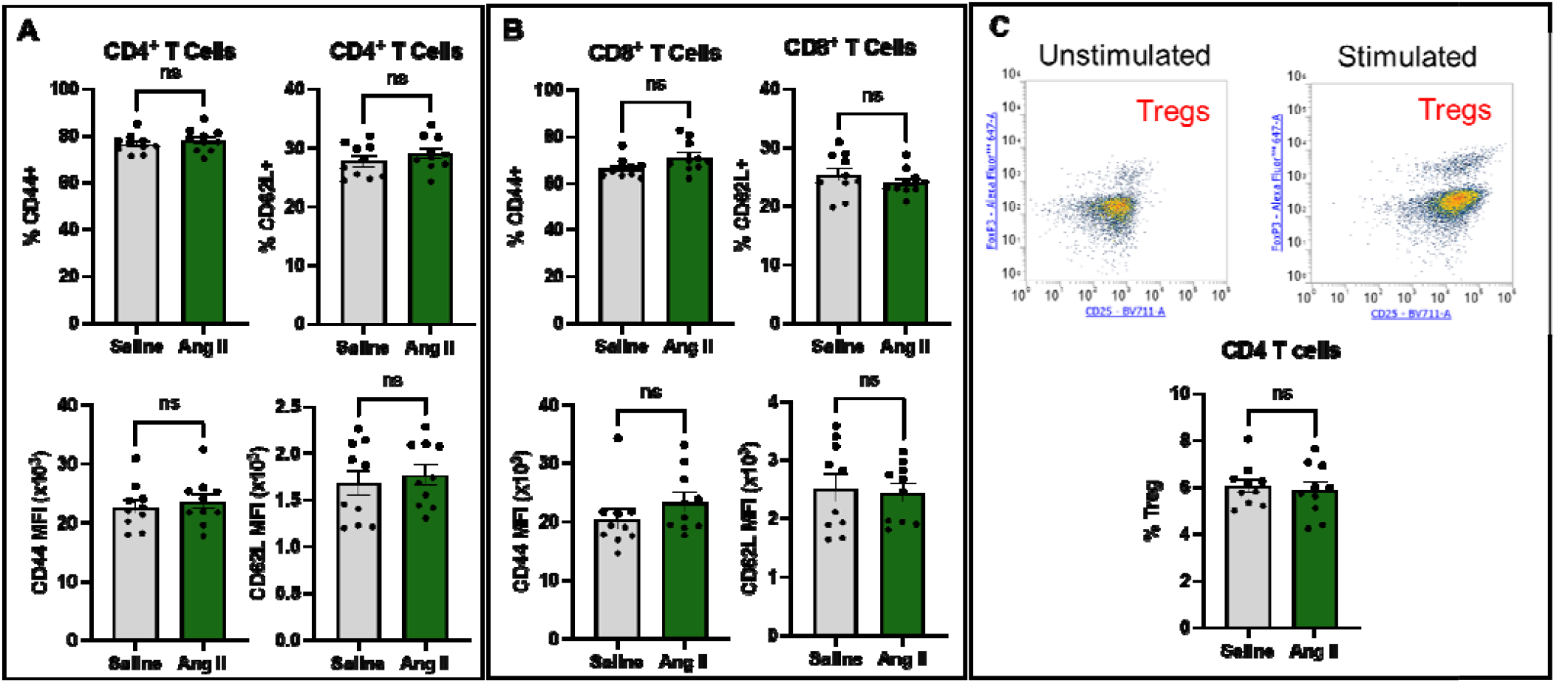
Effects of Ang II on T cell activation markers. T cell activation markers in A) CD4^+^ and B) CD8^+^ T cells, and C) percentages of T regulatory (Treg; CD4^+^ CD25^+^ FoxP3^+^) populations 24 h post-stimulation (n=9-10 mice/croup). Results are depicted as mean ± SEM. Statistical analysis was conducted using unpaired two-tailed t-teste (Welch’s correction used for comparing data of unequal variance) (a=0.05)..

### Ang II promotes induction of cytokines associated with type II and type III interferon responses

To determine the impact of Ang II on cytokine secretion in activated immune cells, we collected cell supernatants 24 and 120 h post activation with anti-CD3/anti-CD28. The increase in the expression of the early activation markers CD69 and CD25 did not correlate with a global increase in cytokine production in the Ang II group. Rather, Ang II caused a selective increase in IFNγ (at 120 h), IP-10 and IL-28B (IFNλ3), but the majority of cytokines were not statistically different between saline and Ang II groups (Fig. 4). In contrast to the increase in IFNγ, IP-10 and IFNλ3, Ang II also caused a decrease in IL-23 secretion. Notably, there were commonalities among the cytokines increased by Ang II in that they were associated with either type II or type III interferon responses both of which play a role in antiviral immunity. There were other cytokines that were not significantly different but were also trending towards an increase. At 120 hours, these included: IL-4 (p=0.065), MIG (p=0.051), MIP-1b (p=0.051), RANTES (p=0.057), VEGF (p=0.162) (Fig. 5 and 6). Overall, these results demonstrate that Ang II promotes a selective increase in certain cytokines associated with type II and type III interferon responses but does not significantly modulate the expression of most cytokines induced in this model.

**Figure 4.**
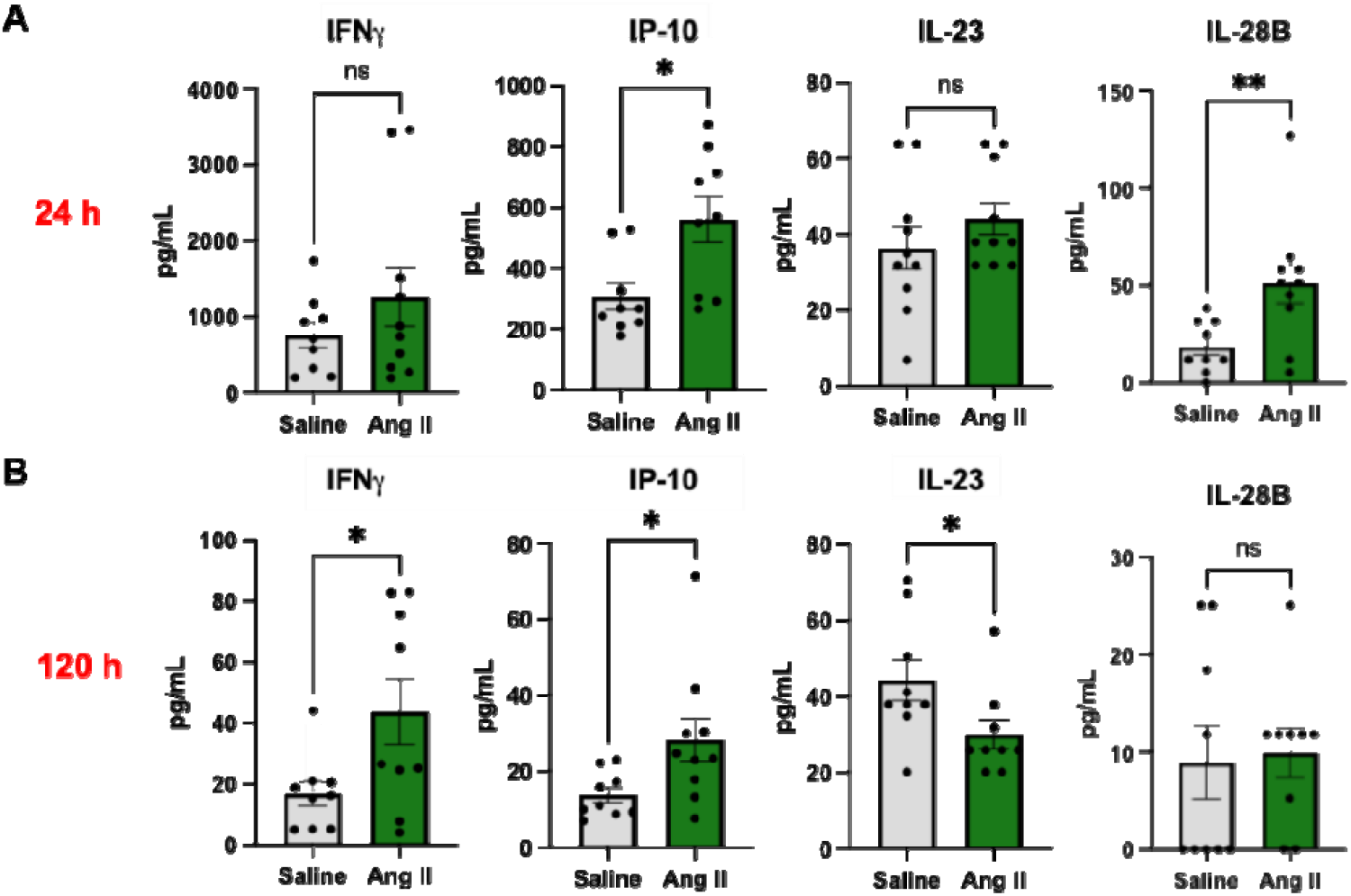
Effect of Ang II on cytokine Induction. Mouse splenocyte cytokine production 24 n and 120 h post-stimulation (n=10 mlce/group). Outliers were determined and removed followed by unpaired two-tailed T-tests (Welch’s correction used when comparing unequal variances and Mann-Whitney test was used for data with non-Gaussian distributions). Differences were observed in IP-10 and IL-28B at 24 h. & IFNγ IP-10. & IL-23 at 120 h (*=p<0.05. **=p<0.01).

**Figure 5.**
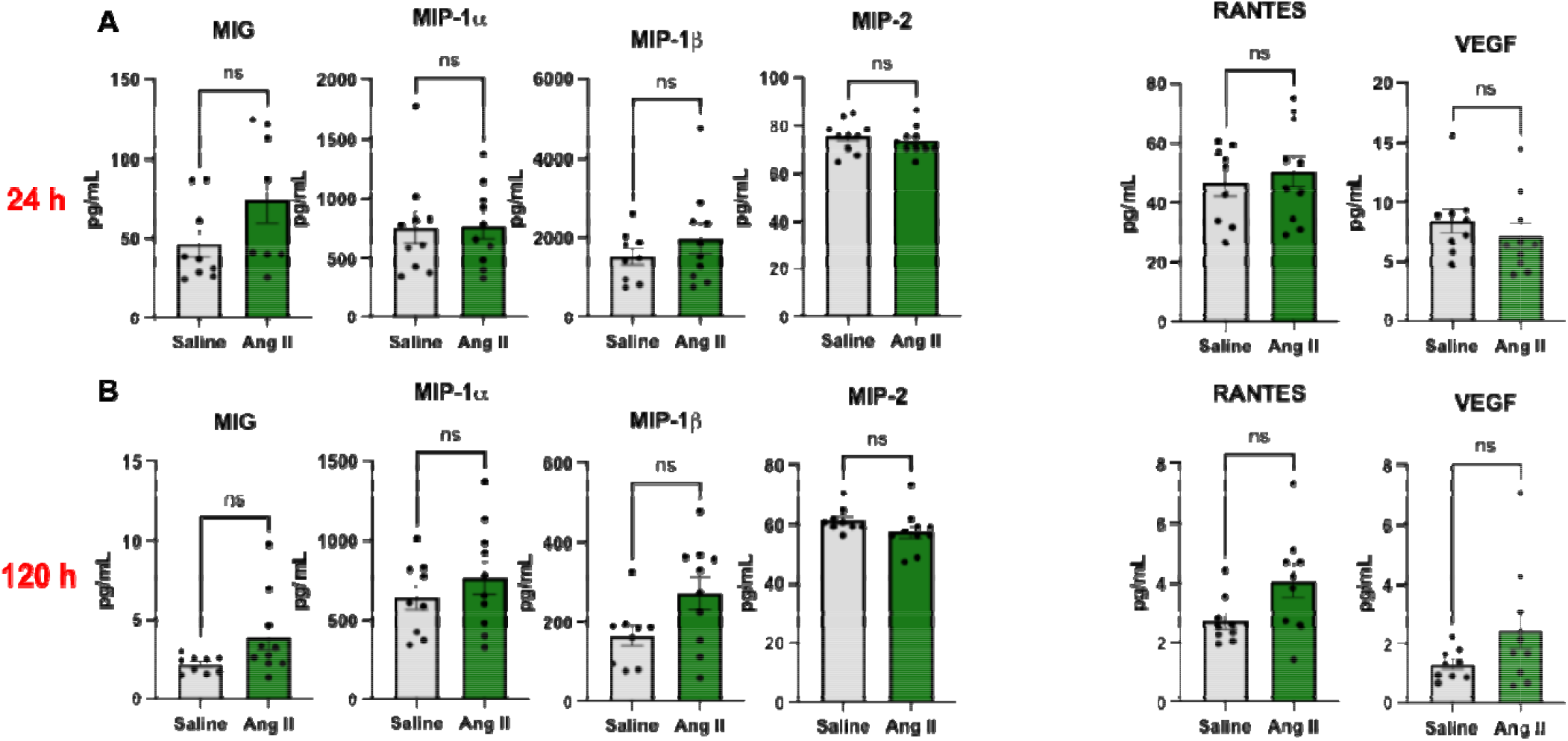
Additional effects of Ang II on cytokine induction. Mouse splenocyte cytokine production 24 (A) & 120 h post-stimulation (r=9-10 mice/group). Outliers were determined & removed followed by unpaired two-tailed T-tests (Welch’s correction used when comparing unequal variances (*c<0.05).

**Figure 6.**
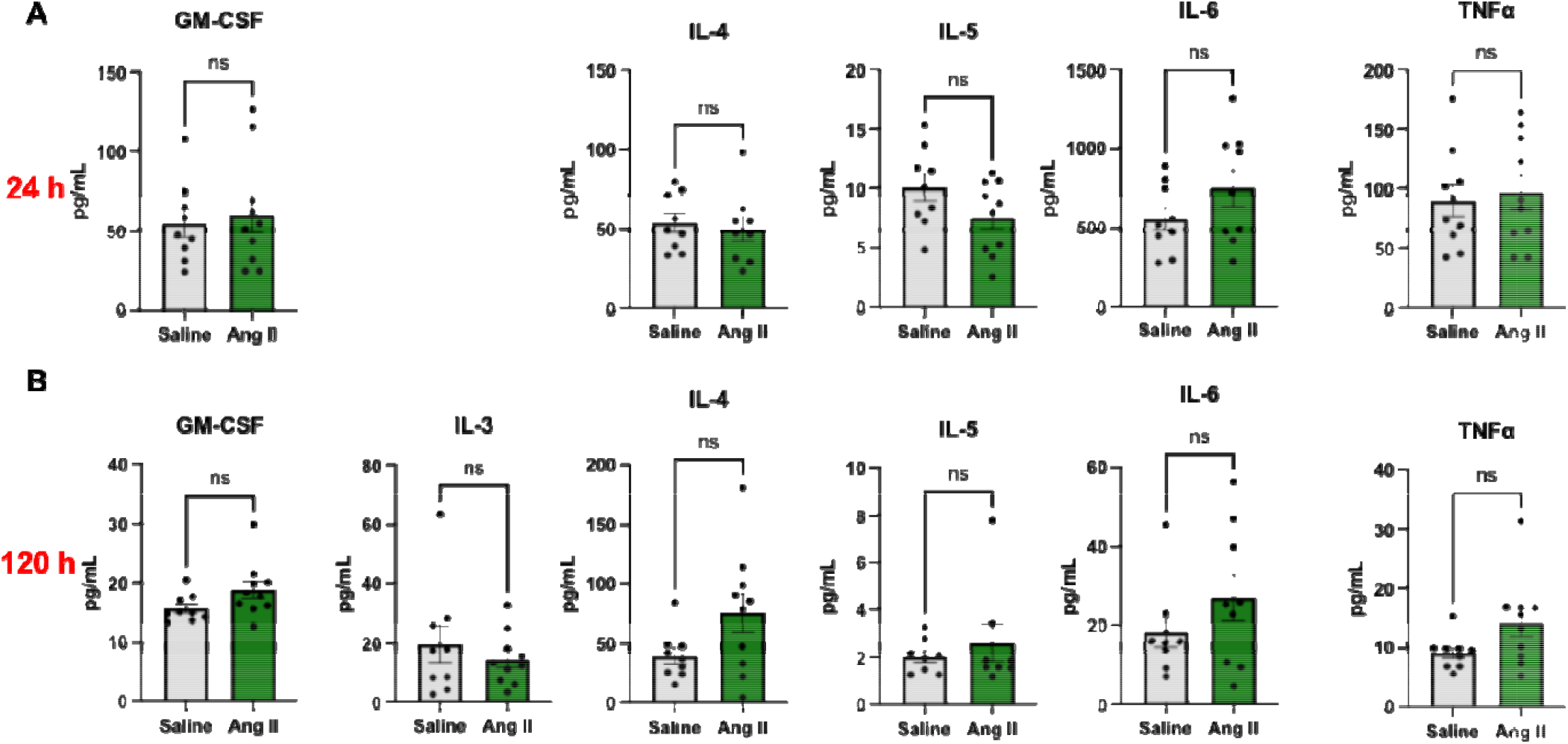
Effect of Ang II on cytokine induction (trending but non-statistically significant cytokines). Mouse splenocyte cytokine production 24 & 120 h post-stimulation (n=9-10 mice/group). Outliers were determined & removed followed by unpaired two-tailed T-tests (Welch’s correction used when comparing unequal variances (a<0.05).

## Discussion

The purpose of this study was to test the hypothesis that Ang II infusion promotes T cell activation and cytokine induction. Toward this end, we activated immune cells isolated from mice following a 2-week infusion with either Ang II or saline. Ang II promotes the induction of activation markers in activated CD8, and to a lesser extent, CD4 T cells. The increased activation marker expression correlated with increased expression of select cytokines (IFNγ, IP-10 and IFNλ3) associated with type II and type III interferon responses, however the majority of cytokines were not statistically different between the saline and Ang II groups. Taken together, the data suggest Ang II may play a role in promoting cytokines associated with an antiviral response.

The cell surface receptors and cytokines induced by Ang II in this study have important immune functions and in some cases are associated with cardiovascular disease. CD137, also known as 4-1BB, is a co-stimulatory molecule expressed on T cells, and CD137 signaling is implicated in atherosclerosis. IL-28B, also known as IFNλ3, is a type III interferon that has been shown to upregulate IFNγ production in NK cells in the lungs and may promote Th1 skewing of CD4 T cells (19, 20). IFNγ is primarily produced by Th1 cells, natural killer (NK) cells, and CD8^+^ cytotoxic T cells, and skews macrophages towards an M1 phenotype. M1 macrophages in turn secrete IP-10, which acts as chemoattractant for T cells. A study that investigated the association of IP-10 and cardiovascular disease in African American populations, assessing data from the Jackson Heart Study (JHS) cohort and the REasons for Geographic and Racial Differences in Stroke (REGARDS) study, found no associations between IP-10 and stroke or coronary heart disease, but did find associations between IP-10 and left ventricular hypertrophy, incident heart failure, and overall mortality (16). Another study found a positive correlation between serum IP-10 levels and blood pressure in 39 patients with essential hypertension (17).

There have been other studies that have linked Ang II to cytokine/chemokine production. With regards to links between Ang II and IP-10, human endothelial cells exposed to Ang II have been shown to have increased IP-10 production through AT_1_R (18). Other groups have indicated that Ang II increases with TNFα production in T cells post-activation. Guzik et al., showed an increase in TNFα in splenocyte supernatants from Ang II-treated mice at 48 hours with stimulation using anti-CD3 (13). Our data show no significant differences in TNFα at 24 hours and 120 hours, however there is a trend towards increased TNFα, along with IL-4, MIP-1b, RANTES, and VEGF in the Ang II group at 120 hours. It is possible that TNFα (along with these other cytokines) would have been significantly different at 48 hours if we had looked at that timepoint. The differences in outcome could also be due to differences in activation protocol in which the current study utilized soluble anti-CD3 and anti-CD28 combined with a cross-linking antibody which results in a moderate level of activation. In contrast, the study by Guzik et al. utilized plate-bound anti-CD3 which is a stronger activator.

The results of this study were somewhat unexpected in the respect that the effects of Ang II in this particular model were much more modest than we had anticipated. While Ang II promoted the expression of activation markers, such as CD69 and CD25, in both CD4 and CD8 T cells, the effects were more modest than we expected. Furthermore, other activation markers, such as CD44 and CD62L (a negative marker that is rapidly downregulated after activation), were not impacted by Ang II at all. Likewise, we observed mixed effects on cytokine induction. While we observed significant induction of IFNγ, IP-10 and IFNλ3, no other cytokines were significantly upregulated by Ang II infusion. The statistically significant effects of Ang II occurred at different time-points. While Ang II did not cause a statistically significant increase in IFNγ at 24 h, there was a non-significant trend that looked similar in magnitude to the effect of Ang II at 120 h. The increased variability of IFNγ at the 24 h time-point likely impacted the statistical analysis. Conversely, Ang II increased IL-28B/ IFNλ3 at 24, but not 120 h after activation. However, IFNλ3 is not directly induced in activated T cells, but rather, is upregulated secondarily due to the effect of T cell cytokines on downstream myeloid or stromal cells. It peaks at early time-points and is likely largely downregulated back to baseline at the 120-h time-point. Thus, Ang II markedly increases IFNλ3 at its peak time-point, but not at 120 h--long after it has returned to baseline levels.

### Limitations

A caveat of this study is that sex-dependent effects of Ang II were not assessed, nor were the underlying mechanisms of the Ang II-mediated increase in activation markers or cytokine secretion determined. Also, the limited number of timepoints (24- and 120-hours) may have been insufficient for capturing significant differences in some cytokines/chemokines, due to the kinetics of the response. Lastly, it is important to consider that this study assessed full splenocyte fractions, which included antigen presenting cells, rather than enriched T cells. Therefore, the extent that Ang II directly affects the response of T cells verses indirectly promoting T cell activation through effects on costimulatory immune cells was not determined in this study. A more thorough characterization of Ang II effects on inflammation is needed to build on these findings that Ang II administration increases T cell propensity to activation.

## Conclusion

Overall, this study demonstrates that Ang II promotes induction of activation markers in CD8, and to a lesser extent, CD4 T cells. The increased expression of activation markers correlates with increased expression of select cytokines associated with type II and type III interferon responses. Taken together, the data indicate that Ang II may contribute to interferon-mediated inflammation and may also play a role in antiviral immunity.

